# Single cell transcriptomics and development of gametocyte-specific molecular markers for avian malaria parasites

**DOI:** 10.1101/2025.07.24.666574

**Authors:** A Berthomieu, S Gandon, AM Talman, A Rivero

**Author notes:** co-last authors (AM Talman, A Rivero).

## Abstract

Avian malaria, caused by *Plasmodium* parasites, poses a significant threat to bird populations worldwide, particularly in vulnerable island ecosystems. Yet, progress in understanding avian malaria transmission dynamics has been hampered by the lack of molecular tools to quantify and sex the transmissible stages of the parasite: the male and female gametocytes. This challenge is compounded by the nucleated erythrocytes of avian hosts and the absence of an *in vitro* culture system, which have historically hindered the advancement of molecular approaches. Here, we develop and validate the first molecular markers to discriminate between asexual, male, and female gametocytes in the widespread avian malaria parasite *Plasmodium relictum*. Using single-cell RNA sequencing and orthology-guided stage mapping, we identified conserved, stage-specific transcripts and leveraged this information to develop molecular markers capable of quantifying and distinguishing male and female gametocytes in two parasite cyt-b lineages, pSGS1 and pDELURB. These markers outperformed microscopy in sensitivity, detected gametocytes earlier and for longer, and revealed consistently higher male-to-female gametocyte ratios than estimated by blood smears. The high degree of conservation of these markers across *Plasmodium* species suggests broad applicability of these markers to other avian lineages.

## Introduction

Although malaria parasites (*Plasmodium* sp.) are largely known for infecting humans, these protozoans can also be found infecting hundreds of other terrestrial vertebrate species, including non-human primates, ungulates, rodents, bats, lizards and birds. Avian parasites are the most widespread, prevalent and diverse of all malaria parasites known to date (Rivero & Gandon, 2018). Transmitted by mosquitoes, avian *Plasmodium* parasites pose a major threat to bird biodiversity and have caused population declines and extinctions, most notably among native Hawaiian species (Atkinson et al., 2014; Garamszegi, 2011). The ongoing impacts of global warming are exacerbating the invasive potential of these parasites, expanding the geographic and ecological range of their vectors and increasing the vulnerability of endemic bird populations . For instance, *Plasmodium* infections are imperiling endemic bird species in the Galapagos Islands and New Zealand, underscoring the urgent need for effective tools to study and mitigate avian malaria (Miranda Paez et al., 2022).

The life cycle of avian *Plasmodium* parasites is similar to that of its human and rodent counterparts (Sinden, 2015). Within red blood cells, the haploid parasite undergoes multiple rounds of asexual multiplication before irreversibly differentiating into sexual stages, called *gametocytes*. Male and female gametocytes are the only stages capable of being transmitted to a mosquito vector. Once in the vector, male gametocytes undergo mitosis to produce eight flagellated male gametes, while female gametocytes mature into single female gametes. Male and female gametes fuse in the mosquito midgut to produce a zygote, which continues the parasite’s cycle inside the mosquito, eventually producing sporozoites transmissible to a new bird in its salivary glands (Sinden, 2015).

The timing and quantity of gametocyte production, along with the sex ratio of gametocytes, are critical epidemiological parameters as they directly influence the success of transmission to the mosquito vector. The ability of the asexual stage to modulate its conversion rate into gametocytes is widely interpreted as a form of adaptive phenotypic plasticity, allowing the parasite to make dynamic life-history adjustments and optimize its transmission strategy in response to environmental cues. Research on human and rodent malaria parasites has indeed shown that *Plasmodium* can adjust both the number and sex ratio of gametocytes in response to a hostile host environment, such as when the host is anaemic (Cameron et al., 2013; Reece et al., 2005) or exposed to antimalarial drugs (Buckling et al., 1999; Schneider et al., 2018; Talman et al., 2004). Other work has shown that gametocyte sex ratios become more male biased in mixed-genotype infections (Reece et al., 2008a) or to ensure female fertilization in the mosquito when gametocytaemias are low (Mitri et al., 2009).

Being able to quantify and sex gametocytes is therefore essential for elucidating both the epidemiology and evolutionary biology of avian *Plasmodium*. Blood smear microscopy is a standard method for detecting malaria parasites, but it has several inherent limitations. Its detection threshold is influenced by factors such as smear quality, the expertise of the examiner, counting method, and the time dedicated to the examination. While examining a larger number of cells and extending the observation period can improve the detection of gametocytes, routine microscopy often underestimates gametocyte prevalence and provides inaccurate counts. Male and female gametocytes can exhibit subtle morphological differences, which are not always distinct from each other or from asexual stages transitioning to gametocytes (Valkiūnas, 2005). Misidentification of parasite stages or host cell artifacts can therefore lead to inaccurate estimates of the gametocyte sex ratio. These issues underscore the need for molecular tools to quantify and sex gametocytes in avian *Plasmodium*.

Unlike the chromosomal or genetic sex determination systems found in many other organisms, *Plasmodium* relies on a distinct molecular mechanism to regulate sexual differentiation. In human and rodent *Plasmodium* species sexual differentiation begins when intraerythrocytic parasites commit to a sexual fate, a process regulated by the transcription factor AP2-G (Kafsack et al., 2014; Sinha et al., 2014). Following this initial commitment, the sexually destined progenitor parasites make a secondary decision to differentiate into either male or female gametocytes (Gomes et al., 2022). Whilst there have been molecular tools available for quantifying and sexing gametocytes in human in rodent malaria for almost two decades (Babiker & Schneider, 2008) the development of comparable molecular tools for avian *Plasmodium* has been hindered by several significant challenges. Firstly, the nucleated nature of avian erythrocytes results in an overwhelming abundance of host nucleic acids, which obscures and complicates the detection and quantification of parasite-specific genetic material. Secondly, the asynchronous development of the parasite means there is no specific time point when only gametocytes are present in the blood. Finally, no *in vitro* culture systems exist for avian *Plasmodium*, making it difficult to isolate and study gametocyte-specific stages under controlled conditions.

In this study, we present the first single-cell (SC) transcriptome analysis of an avian malaria parasite and leverage it to develop and test molecular markers for quantifying and sexing gametocytes in *Plasmodium relictum,* one of the most widespread and prevalent species within the avian *Plasmodium* genus (Rivero & Gandon, 2018). The single-cell transcriptome provided a global view of gene expression, allowing us to apply a "guilt-by-association" approach by comparing expression patterns with those documented in the Malaria Cell Atlas (Dogga et al., 2024; Howick et al., 2019). We selected two stage-specific markers for female gametocytes, male gametocytes, and asexual stages, and validated these markers in an experiment using two different *P. relictum* cyt-b lineages (pSGS1 and pDELURB4). These markers represent a significant step forward in our ability to study avian malaria at a molecular level, providing a foundation for future research into the epidemiology and evolutionary dynamics of host-parasite interactions under changing environmental conditions.

## Materials and Methods

### Single cell transcriptomics and stage-specific marker identification

We adapted the protocol of Reid *et al*. (2018) originally developed for *Plasmodium berghei* and *Plasmodium falciparum* to perform single-cell transcriptomics on canary blood containing both sexual and asexual stages of *Plasmodium relictum*. For this purpose, we infected one canary (*Serinus canaria*) with *Plasmodium relictum* (lineage pSGS1) through intraperitoneal infection using 100 µl of infected blood from our infected canary stock. The bird was sampled 10 days post infection, and its parasitaemia and gametocytaemia were quantified via the microscopic examination of blood smears (4% and 0.9%, respectively). Fourty microliters of the sampled blood were placed in a 1.5ml Eppendorf tube, and the nuclear and mitochondrial DNA were stained by adding 400 µl of a mixture of SYBR green (1/4000) and MitoTracker-DeepRed (1/2000) in phosphate-buffered saline (PBS). The mixture was kept on ice for 15 minutes and centrifuged for 1 minute at 2000 g (all centrifugation steps were performed at 4°C). The supernatant was discarded and the pellet was resuspended in 400 µl PBS containing 0.15% saponin to lyse the red blood cells for 5 minutes on ice. After a 2 min centrifugation at 2000 g, the supernatant was discarded and the pellet was washed twice with 1ml of PBS followed by 2 min centrifugation at 2000g. Finally, the supernatant was discarded, the pellet was re-suspended in 500 µl of PBS, and the parasite suspension was kept on ice in the dark before the subsequent flow cytometry step.

The Sybr green/Mitotracker-stained samples were sorted with a FACS ARIA IIIU BECTON DICKINSON cell sorter into a 96-well RNAse-free plate (Abgene). Each well of the plate contained a mix of 2 µl of lysis buffer (0.8% Triton-X in nuclease-free water), 1µl of 10 mM dNTPs, 0.1µl of 10µM OligoDT 30 (5’-AAGCAGTGGTATCAACGCAGAG TAC(Tx30)3’), 0.1µl of 20U SuperRNAsin (Life Technologies) and 0.8 µl of nuclease-free water (Ambion). The sorting gate for *P. relictum* infected blood had been previously established by comparing infected versus non-infected blood (Figure S1A). Parasites were sorted by gating single-cell events, with controls including 3 wells containing 10 events from the infected sample and a non-sorted negative control well. Sorted plates were spun at 1000 rpm for 10 s and immediately placed on dry ice.

Reverse transcription was carried out according to the protocol of Reid *et al*. (2018). Briefly, the sorted plates were incubated at 72°C for 3 min before adding 5.7µl of the reverse transcription master mix (Table 1). The plates were then incubated using the following programme: 1 cycle at 42°C for 90 min; 10 cycles of 42°C for 2 min and 50°C for 2 min) and 1 cycle at 70°C for 15 min. After incubation, 2.5 µl of 2X KAPA Hotstart HiFi Readymix and 0.25 µl of a 100 µM ISO SMART primer solution (5’-AAGCAGTGGTATCAACGCAGAGT-3’, Picelli et al., 2013) were added to each well. The plates were re-incubated using the following cycling programme: 1 cycle at 98°C for 3 min, followed by 30 cycles of 98°C for 20 s, 67°C for 15 s and 72°C for 6 min, ending with 1 cycle at 72°C for 5 min. Samples were then purified with 1X Agencourt Ampure beads (Beckman Coulter) in a Zephyr G3 SPE Workstation (Per-kin Elmer) according to the manufacturer’s instructions. The amplified cDNA was eluted in 10 µL nuclease-free water. The quality of synthesised cDNA was monitored with the high-sensitivity DNA chip on an Agilent 2100 Bioanalyser on 10 randomly selected cDNA samples. Libraries were prepared using the Nextera XT kit (Illumina) according to the manufacturer’s recommendations, using 96 different index A combinations. After indexing, libraries were pooled for clean-up in a 4:5 ratio of Agencourt Ampure beads (Beckman Coulter). The quality of the libraries was monitored with the high sensitivity DNA chip on an Agilent 2100 Bioanalyser. Empty-well controls and single cells were pooled separately from 100-cell controls and loaded proportionally to their expected cell content. The 96 libraries were sequenced on a MiniSeq (Illumina) with 50PE and 1% PhiX. The demultiplexed sequencing data was processed as follows: FASTQ files were trimmed using trim_galore (v0.4.1) with the parameters -q 20 -a CTGTCTCTTATACACATCT --paired -- stringency 3 --length 50 -e 0.1. HISAT2 (v2.0.0-beta) (Kim et al., 2019) indexes were generated for the *Plasmodium relictum* (SGS1) from GeneDB (Logan-Klumpler et al., 2012) in March 2020, using default parameters. Trimmed paired-end reads were aligned to the respective genome using hisat2 --max-intronlen 5000 -p 12. Gene annotation files were downloaded from GeneDB (March 2020). When multiple transcripts were annotated for a single gene, only the primary transcript was considered. Read counts were assigned to genes using HTSeq (htseq-count -f bam -r pos -s no -t CDS, v0.6.0 ) (Anders et al., 2015). By default, HTSeq excludes multimapping reads (-a 10), ensuring that ambiguously mapped reads from gene families with similar sequences were not included in the analysis. The raw sequence counts are provided in file SX.The dataset was processed using single cell experiment (v1.26.0) and scater (v1.32.1) (McCarthy et al., 2017). Genes with more than 5 reads in at least two cells were retained for subsequent analysis. Control wells ase well as cells with fewer than or equal to 10,000 total read counts and/or fewer than 400 genes were filtered out. After filtering, the *P. relictum* dataset retained 63 out of 92 single sorted cells, detecting 4,117 unique genes.

**Table 1.**
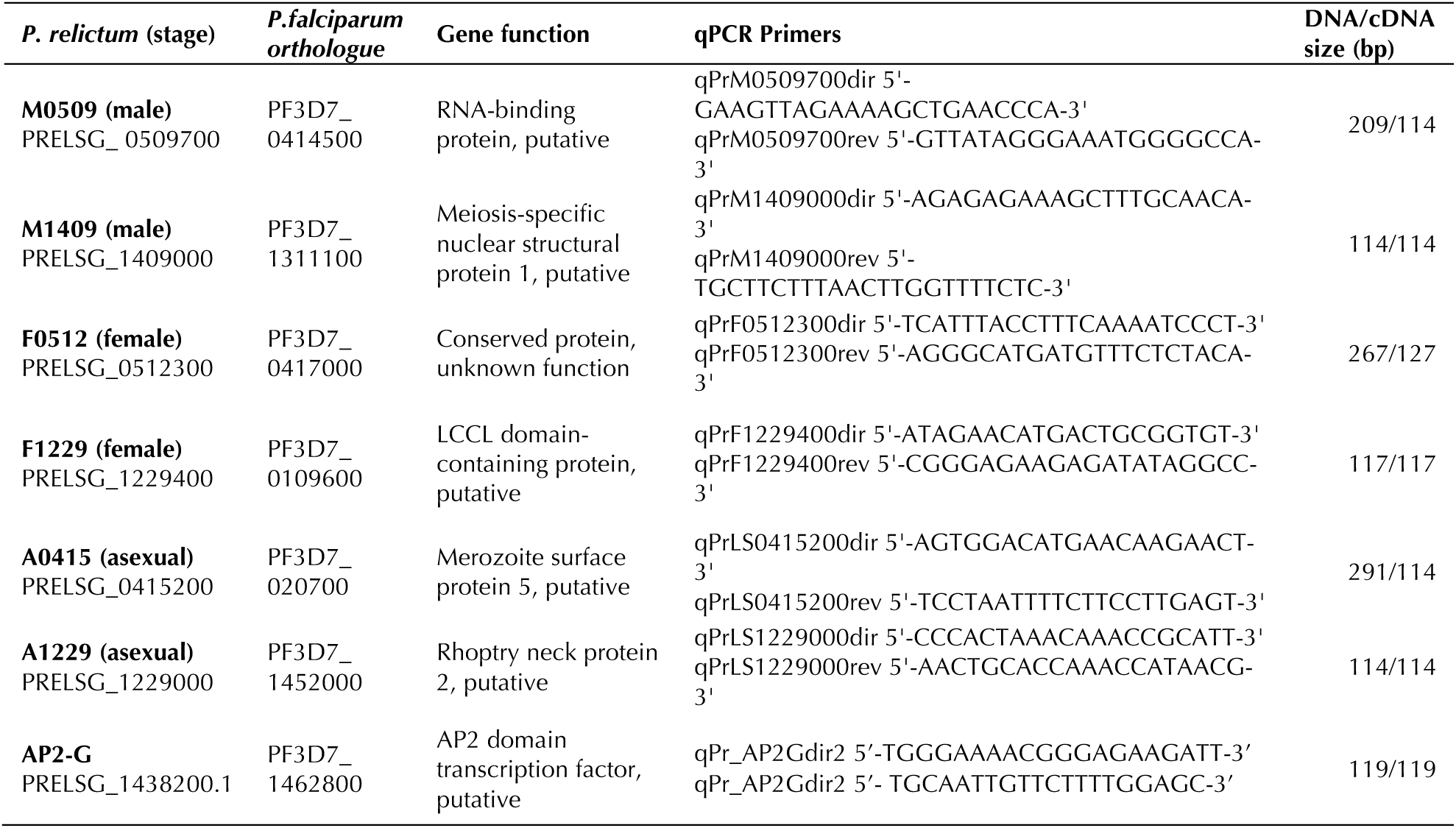
Stage-specific genes *Plasmodium relictum* (male, female, asexual), the molecular marker for conversion (AP2-G), and the corresponding primers.

Dimentionality reduction was perfomed within scater (v1.32.1) (McCarthy et al., 2017). We estimated the k for clustering using SC3 *sc3_estimate_k* function and performed clustering and marker identification with the same package. Stage mapping was performed by transforming the data by orthology to *P. berghei* using the OrthoMCL data (Chen et al., 2006) (March 2020) and selecting only 1:1 orthologs. The transformed data was then mapped onto a reference *P. berghei* dataset (Russell et al., 2023) using scmap (Kiselev et al., 2018) with a similarity threshold of 0.4. Dowstream analyses were greatly facilitated by using PlasmoDB (Amos et al., 2022).

We selected and designed primers for amplifying fragments of two female gametocyte, two male gametocyte, and two asexual stage genes (**Table 1**). Primers were designed with *P. relictum* lineages pSGS1 and pDELURB4 sequences using Primer 3 software (version 4.1.0). Whenever possible, primer pairs were placed on either side of an intron to ensure the amplified fragment was derived from cDNA and not from contaminating genomic DNA.

### Experimental validation of stage-specific markers

To validate the stage-specific markers, we designed and conducted an experiment aimed at testing their specificity and accuracy using bulk RNA transcriptomics. Birds were infected by intraperitoneal injection of 100 µl blood infected with either pSGS1 or pDELURB4 (3 birds each). The blood of these experimental birds was sampled on days 0, 3, 5, 7, 10, 12, 14, 17, 19, 21, 24 and 31 post infection. At each sampling event, 10 µl of blood was mixed with 100µl of PBS and stored at -20°C while an additional 10 µl was mixed with 500µl of Trizol Reagent (Invitrogen) and stored at -80°C for subsequent DNA and RNA extraction, respectively. DNA was used for molecular quantification of the total parasite load, while RNA was used to quantify the different stages. In addition, blood smears were prepared for the microscopic assessment of parasitaemia and gametocytaemia, providing an additional method of quantification alongside the molecular marker analysis. DNA and RNA were extracted with the Blood and Tissue kit (Qiagen) or with Trizol (Invitrogen), respectively, following the manufacturer’s instructions. DNA was eluted in 80 µl of Nuclease-free water (Qiagen) and the RNA pellet was resuspended in 40 µl of DEPC-treated water (Invitrogen). Both of DNA and RNA concentrations were measured using a Spark spectrophotometer (Tecan).

Reverse transcription was conducted on the RNA samples using SuperScript II RT (Invitrogen) following the manufacturer’s instructions with a few minor modifications. To this end, 100 ng of RNA was mixed with 0.5 pg of oligo(dT) (Invitrogen), 1 µl of random decamers (Invitrogen), 1µl of 10mM each dNTP mix and completed to 12 µl with nuclease-free water (Qiagen). The mixture was heated at 65°C for 5 min, then quickly chilled on ice. Next, 4µl of 5X First-Strand Buffer, 2µl of 0.1M DTT and 40U of RNase OUT (Invitrogen) were added to each well. The plate was incubated at 42°C for 2min, followed by 25°C for 2 min. Finally, 200U of SuperScript II RT was gently added to each well, and the plate was incubated first at 25°C for 10 min, then at 42°C for 50 min. The reaction was inactivated by heating at 70°C for 15 min.

For each bird, stage-specific quantification was performed using qPCR on a LightCycler 480 (ROCHE) in a single run. Reactions were conducted in a final volume of 6 µl, containing 1 µl of cDNA, 0.6 µM of the corresponding primer pair (**Supplementary Table S1**), and 3 µl of SensiFAST SYBR No-ROX kit mix (OZYME). Amplifications were conducted in triplicate, using the following cycle: 8 min at 95°C, followed by 45 cycles (4 s at 95°C, 20 s at 62°C, 20 s at 72°C), and ending with the melting curve program (70-98°C). When the primer pairs were designed on either side of an intron, we checked the specificity of the amplicon by monitoring the melting curve of the amplified fragment. Callibration curves were established using a synthetic plasmid (Eurofins) containing a single copy of each target gene. For the quantification of male and asexual stages, the plasmid contained one copy each of the two gene fragments of interest (**Table 1**) and one GAPDH copy (housekeeping gene). For the quantification of female stages, due to the complexity of the gene fragment sequences it was not possible to synthesise a single plasmid. Consequently, two plasmids each containing a single copy of one of the markers plus a copy of GAPDH were synthethised.

Molecular quantification of the total parasitemia was carried out using 2 qPCRs in a single run. The first one amplified a fragment of the *Plasmodium cyt b* gene with the primers L9 (5‘- AAACAATTCCTAACAAAACAGC-3’) and NewR (5’-ACATCCAATCCATAATAAAGCA-3’) from Knowles *et al* 2009 , and the second one amplifyed the bird’s 18s rDNA gene with the primers 18sAv7 (5’-GAAACTCGCAATGGCTCATTAAATC-3’) and 18sAv8 (5’- TATTAGCTCTAGAATTACCACAGTTATCCA-3’) (Cellier-Holzem et al., 2010). Each qPCR reaction was performed in triplicate for each sample, in a final volume of 6 µl containing 5ng of DNA, 0.6 µM of each primer, 3µl of SensiFAST SYBR No-ROX kit mix (Ozyme) on a LightCycler 480 (Roche). The amplification cycle followed the same conditions as previously described.

Expression of the AP2-G transcription factor, which regulates differentiation from asexual to sexual stages, was quantified by qPCR. qPCR primers (**Table 1**) were designed based on the homology of the orthologous *P. falciparum* gene (*PF3D7_1222600*, PlasmoDB). Reactions were performed in triplicate for each sample in a final volume of 6 µl, consisting of 1 µl of cDNA, 0.6 µM of each primer, and 3 µl of SensiFAST SYBR No-ROX kit mix (Ozyme), using a LightCycler 480 (Roche).

### Statistical analyses

To analyze the experimental validation of stage-specific markers we used the R statistical package (v4.1.1). To account for repeated measures within each bird, we constructed mixed models with bird identity as a random effect (all models are summarized in **Supplementary Table S2**). Model selection followed a stepwise approach: starting with a maximal model that included all fixed effects and their higher-order interactions, we sequentially removed non-significant terms to obtain a minimal model. The significance of the explanatory variables was assessed using likelihood ratio tests (LRT) which follow an approximate chi-square distribution (Bolker, 2008), with a significance threshold of p = 0.05. Proportion data (sex ratio) was analyzed using a binomial distribution. Least square means were estimated using the emmeans function. When appropriate, we calculated the marginal R^2^ (the proportion of the variance explained by the fixed effects) following the method of Nakagawa & Shielzeth (Nakagawa & Schielzeth, 2013).

### Ethical statement

Bird manipulations were carried out in strict accordance with the “National Charter on the Ethics of Animal Experimentation” of the French Government. Experiments were approved by the Ethical Committee for Animal Experimentation established by the authors’ institution (CNRS) under the auspices of the French Ministry of Education and Research (permit number CEEA- LR-221 1051).

## Results

### Single cell transcriptomics and stage-specific marker identification

*P. relictum* erythrocytic stages reside within nucleated red blood cells posing unique challenges for isolating infected cells and generating transcriptomes without overwhelming host contamination. To address this, we optimised a host lysis protocol that enables the cell sorting of extra-cellular but intact *P. relictum* erythrocytic stages (**Figure S1A**). Following cell sorting, we performed single cell RNAseq and, after quality control, recovered 63 high-quality parasite transcriptomes. Clustering analysis grouped the cells into five distinct clusters, each characterized by differences in the number of expressed genes per cell (**Figure S1B**). Stage-specific variations in expressed gene numbers per cell have been previously observed in *Plasmodium* single-cell transcriptomes (Howick et al., 2019). Cluster 2 and 4 expressed orthologs of markers corresponding to established female (P47) and male (HAP2) gametocytes in rodent and human malaria parasites (Howick et al., 2019), whereas clusters 3 and 5 were characterized by the expression of the MSP1 ortholog, indicative of maturing asexual stages (**Figure 1A**). To formally assign developmental stages to these clusters, we transformed the data by orthology to *P. berghei* and performed cell mapping onto a reference *P. berghei* dataset (Russell et al., 2023) (**Figure S1C**). All cells from cluster 2 mapped to female gametocytes, while all cells from cluster 4 mapped to male gametocytes. Clusters 1, 3 and 5 mapped to asexual stages.

**Figure 1.**
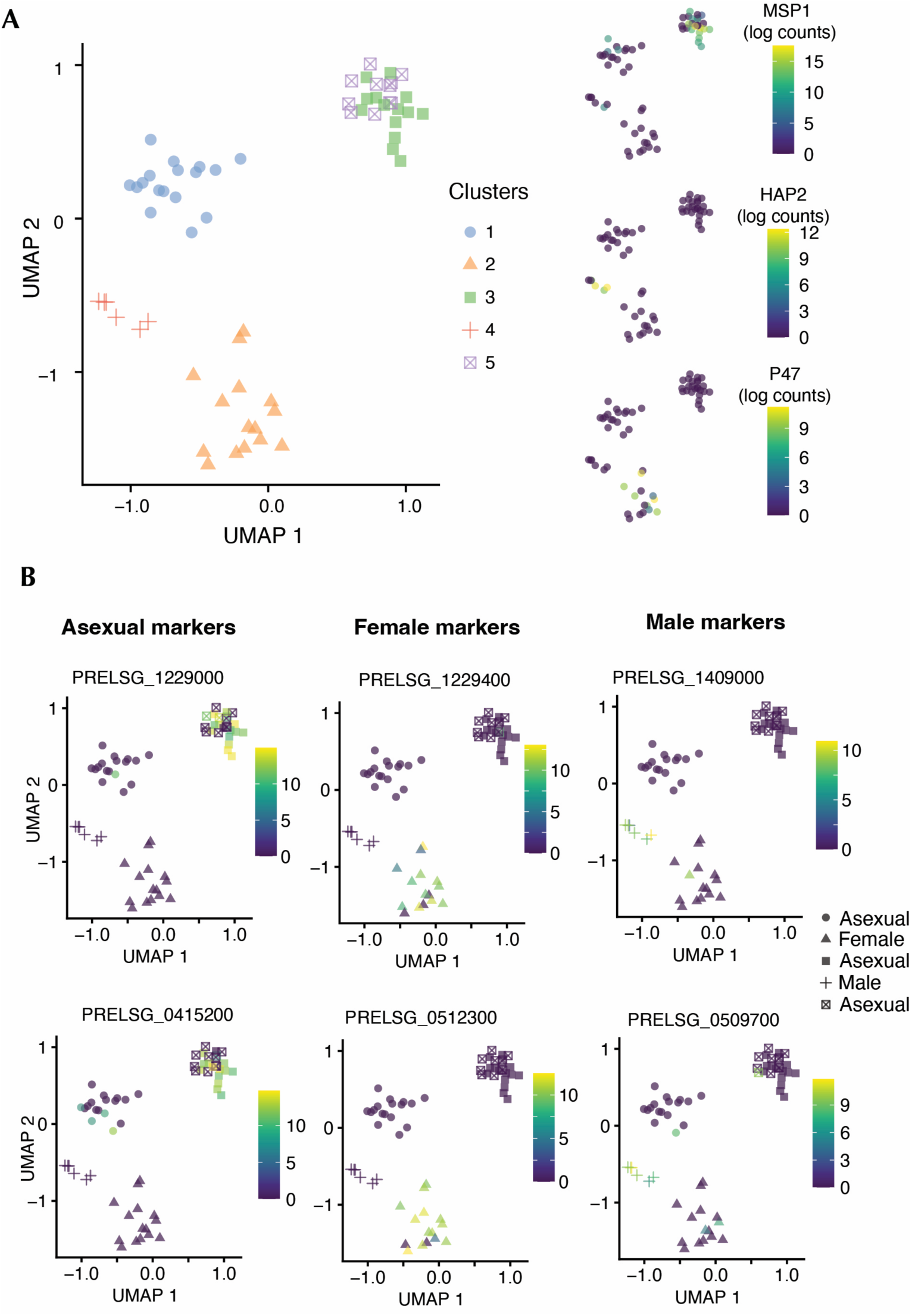
**(A)** UMAP visualization of 63 *P. relictum* transcriptomes, clustered using k-means (left). The expression profiles of orthologs of known female (P47), male (HAP2), and mature asexual (MSP1) parasites in *P. berghei* and *P. falciparum* (right) (PMID: 31439762). **(B)** Expression of *P. relictum* stage-specific markers within each of the five different clusters.

Together, these data provide reference transcriptomes for asexual, male and female gametocytes of *P. relictum*. We further identified stage-specific marker genes for each cluster and selected two robust candidate biomarkers of asexual, female and male parasites (**Figure 1B**). The orthologs of the selected gametocyte markers were also markers of the same stage in *P. berghei* (Russell et al., 2023) and *P. falciparum* (Gomes et al., 2022).

### Experimental validation of stage-specific markers

We assessed the performance of male and female gametocyte markers in two different ways. First, by regressing the number of male and female gametocytes as estimated by the markers against the number of male and female gametocytes observed in the blood smears by microscopy (**Figure 2A**). Statistical analyses revealed a very significant correlation between blood counts and the number of marker copies, although the estimated slope of the relationship was steeper for pDELURB than for pSGS1 (**Supplementary Table S2**, model #1, interaction blood counts: *Plasmodium* lineage *χ*^2^_1_= 29.91, p < 0.0001, estimated marginal means and standard errors: pDELURB = 0.811 ± 0.049, pSGS1: 0.563 ± 0.049, marginal R^2^ = 0.61). This difference between the lineages was independent of the type of marker (model #1, 3-way interaction *χ*^2^_3_= 4.10, p = 0.2506). Using the newly developed markers, we were able to detect gametocytes in 32% of the pSGS1 samples and 21% of the pDELURB samples that were negative for gametocytes through microscopic examination of the blood. Second, by regressing the number of male and female gametocytes as estimated by the markers against the number of copies of the AP2-G gene, a conserved gene responsible for the sexual commitment of *Plasmodium* parasites in blood (**Figure 2B**). Statistical analyses revealed a very significant correlation between AP2-G counts and the number of marker copies (model #2, AP2-G effect *χ*^2^_1_ = 158.01, p<0.0001, marginal R^2^ = 0.60), which was independent of the lineage (model #2, interaction AP2-G: lineage, *χ*^2^_3_ = 3.519, p = 0.318). The markers were less effective at detecting gametocytes than AP2-G. Specifically 31% of pSGS1 and 30% of pDELURB samples that tested positive for gametocytes based on AP2-G expression were not detected as gametocyte-positive using the new markers. Male and female molecular markers outperformed the microscopic estimation of male and female gametocytes from blood smears (**Figure 3**). These markers detected gametocytes as early as, or earlier than, blood counts and remained detectable for a longer duration. Two of the markers, the male marker M1409 and the female marker F0512, outperformed their counterparts in both pSGS1 and pDELURB4 by detecting gametocytes for a longer duration following the peak of infection.

**Figure 2.**
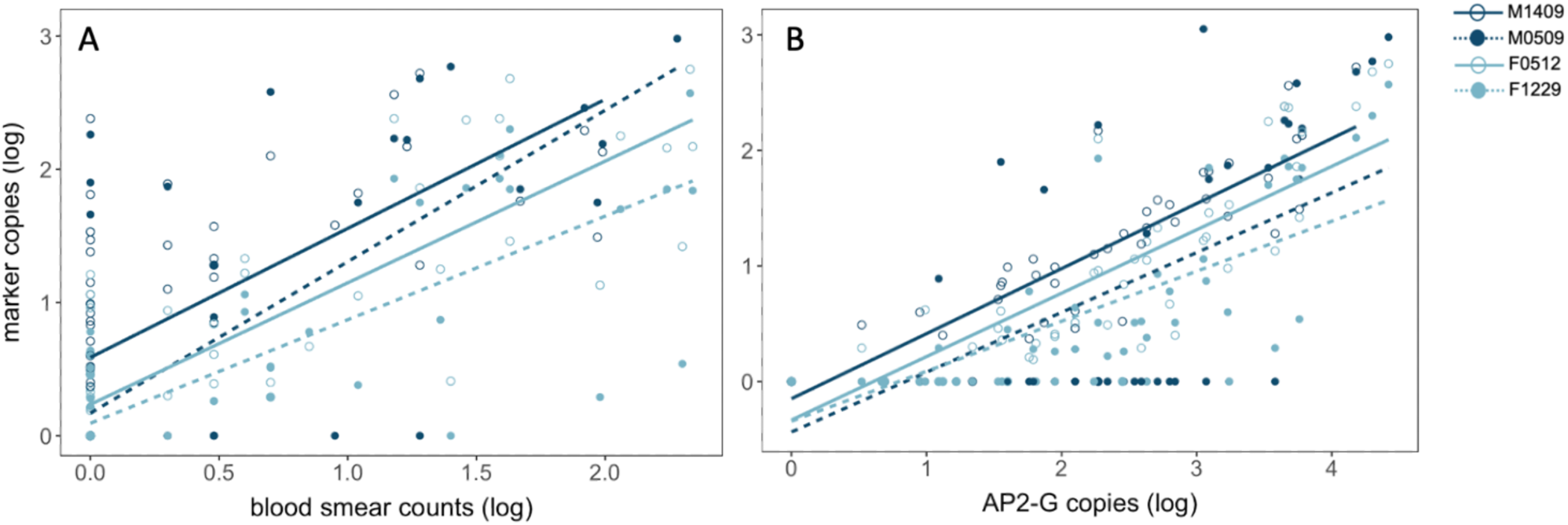
Regression of marker copy number (log-transformed) on blood smear gametocyte counts (A) and AP2-G copy number (B). Male markers (M1409 and M0509) in dark blue, female markers (F0512, F1229) in light blue.

**Figure 3.**
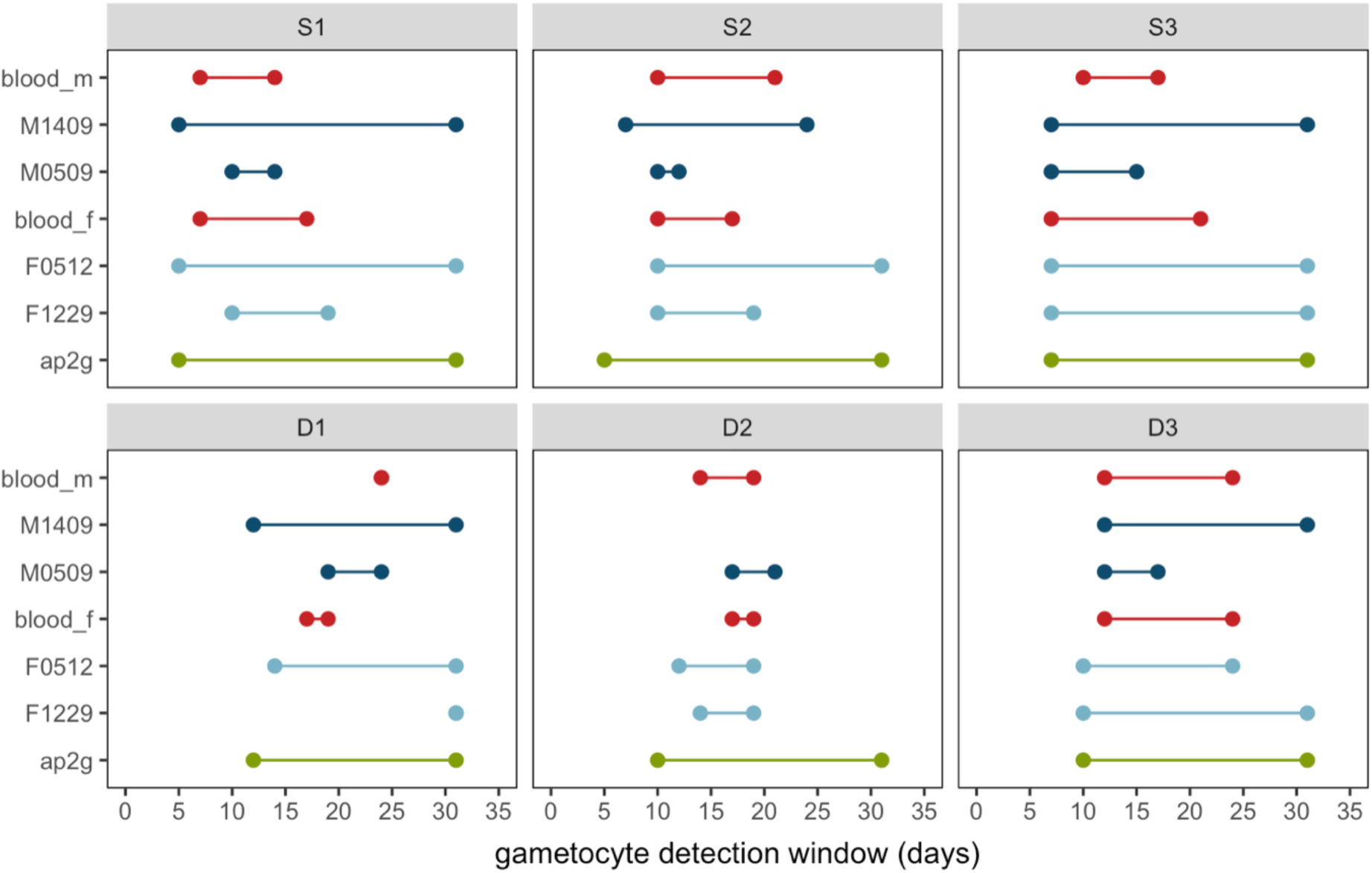
Gametocyte detection window for the different gametocyte quantification techniques: blood smear counts (males: blood_m, females: blood_f), sex-specific molecular markers (males: M1409, M0509, females: F0512, F1229) and gametocyte conversion molecular markers (ap2g). The figure shows the results in birds infected with pSGS1 (S1-S3) and pDELURB4 (D1-D3).

The temporal dynamics of asexual, male and gametocyte stages in each bird is shown in **Figure 4**. With the exception of one bird infected with DELURB4 (**Figure 4F**), which did not exhibit any particular dynamic pattern, all other birds showed male and female gametocyte dynamics that mirrored those of the asexual stages (**Figure 4A-E**). In pSGS1 the first gametocytes were detected between days 5-7 post infection and peaked on days 10–12, whereas in pDELURB4 gametocytes appeared and peaked later (days 10-14 and days 14–19, respectively).

**Figure 4.**
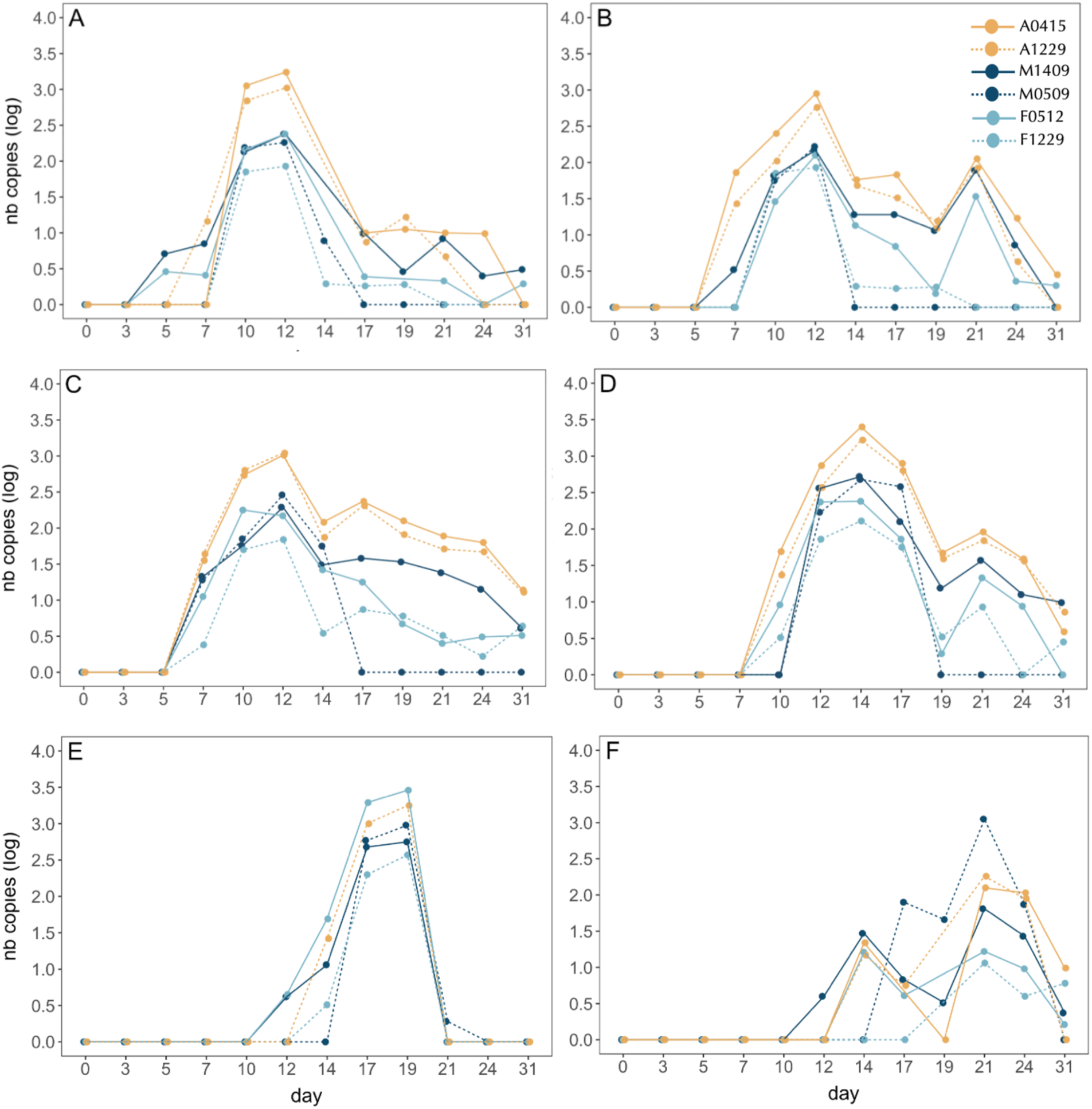
Temporal dynamics of marker copy number (log-transformed) across birds infected with either pSGS1 (A-C) or pDELURB4 (D-F). Asexual markers (A0415, A1229) in yellow, male markers (M1409 and M0509) in dark blue, female markers (F0512, F1229) in light blue.

Overall, molecular markers estimate a higher proportion of male gametocytes compared to blood smears, suggesting a difference in detection efficiency of male and female gametocytes between the methods (**Figure 5**). Consequently, the sex ratio (% males) estimated from molecular markers was significantly higher than that estimated from blood smears (lsmeans = 0.53 and 0.33, respectively, model #3, *χ*^2^_1_ = 5.18, p = 0.022, no difference between the strains *χ*^2^_1_ = 0.002, p = 0.9619).

**Figure 5.**
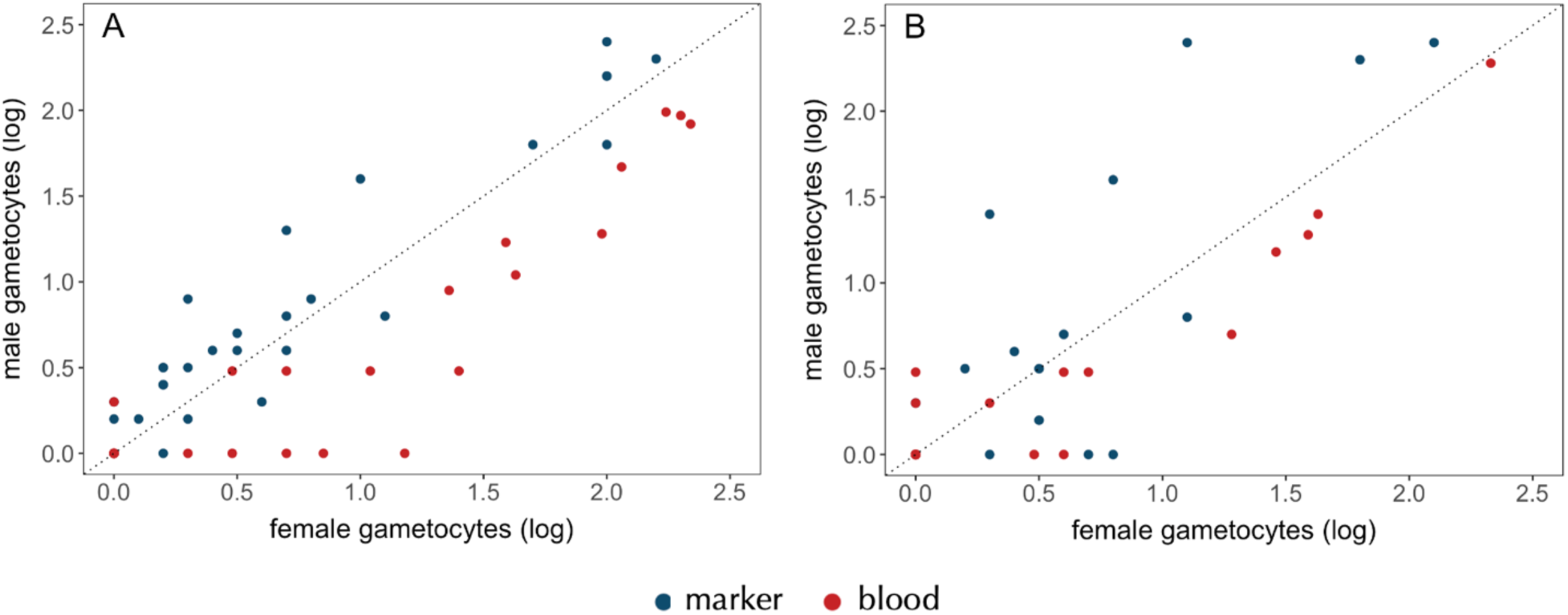
Male and female gametocyte abundance (log-transformed) estimated using molecular markers (blue dots, average of two markers) or blood smear counts (red dots). Data are shown for birds infected with pSGS1 (A) or pDELURB4 (B). The diagonal line (intercept = 0, slope = 1) represents a balanced sex ratio.

## Discussion

Molecular markers to quantify and sex gametocytes have long existed for rodent and human malaria parasites (Babiker et al., 2008), but until now, equivalent tools were lacking for avian systems. In this study, we report the first development and validation of molecular markers for quantifying and sexing gametocytes of *Plasmodium relictum*, a key model in avian malaria research. Our approach, combining single-cell transcriptomics with orthology-guided stage mapping, allowed us to overcome major technical challenges posed by the nucleated red blood cells of avian hosts, and the absence of an in vitro culture system, both of which have traditionally hindered the development of molecular markers for key parasite stages. The host-cell lysis protocol we developed allowed for the isolation and sequencing of *P. relictum* erythrocytic stages with minimal host contamination. This represents a significant methodological advance for avian malaria research and may be applicable to other avian haemosporidian species.

Clustering of the single-cell transcriptomes revealed distinct transcriptional profiles corresponding to asexual stages as well as male and female gametocytes. These assignments were robustly supported by orthology to known stage-specific markers in *P. berghei* and *P. falciparum* (Howick et al., 2019; Russell et al., 2023). The identification of reliable and conserved molecular markers for each stage enabled us to design a set of qPCR assays capable of separately quantifying male and female gametocytes. Two of the markers in particular, the male marker M1409 and the female marker F0512 outperformed the others in terms of sensitivity in the two lineages tested: pSGS1 and pDELURB. These markers consistently detected gametocytes earlier and remained detectable for a longer duration following the peak of infection, making them especially valuable for tracking transmission-relevant stages over time. pSGS1 and pDELURB are phylogenetically close and represent but a small fraction of the over 600 cyt-b lineages described thus far for avian *Plasmodium* parasites (Bensch et al., 2009). Further validation is needed to determine whether these markers reliably capture gametocyte dynamics in lineages with greater genetic divergence. However, given the high degree of conservation of stage-specific gene expression patterns across *Plasmodium* species (Howick et al., 2019) we suggest that these markers are likely to be broadly applicable across other avian *Plasmodium* lineages.

Our experimental validation confirms that these molecular markers significantly outperform traditional microscopic examination in terms of sensitivity and temporal resolution. Importantly, the markers were able to detect gametocytes in over 30% of the pSGS1 and 20% of the pDELURB4 samples scored as negative by microscopy, particularly in the early and late phases of infection. These results are consistent with previous findings in human malaria that suggest microscopy underestimates gametocyte prevalence (Babiker et al., 2008). The molecular estimates also showed a strong correlation with AP2-G expression, the master regulator of sexual commitment in *P. falciparum and P.berghei* (Kafsack et al., 2014; Sinha et al., 2014). The fact that a proportion of AP2-G-positive samples lacked detectable gametocyte markers suggests the potential presence of early committed stages not yet expressing mature gametocyte markers. Future work should explore additional markers that span the entire differentiation process, from early commitment to fully mature gametocytes.

Interestingly, we observed differences in the temporal dynamics of gametocyte development between *P. relictum* lineages, with pSGS1 producing gametocytes earlier than pDELURB4. While these differences could reflect intrinsic differences in the developmental pace of the lineages, they might also stem from variation in parasite load or host response. Canaries, originally from the Macaronesian Islands (Azores, Madeira, and the Canary Islands), have been bred in captivity in Europe since the 17th century. Today, birds housed in outdoor aviaries frequently become natural hosts for *Plasmodium relictum*, including these two lineages. It remains to be determined whether these developmental differences also manifest in wild birds with a longer co-evolutionary history with the parasite. More systematic investigations will be needed to disentangle these effects and assess their implications for transmission.

In *Plasmodium*, sex ratios are generally expected to be female-biased, as each male gametocyte produces up to eight gametes in the mosquito vector, whereas each female gametocyte yields only a single gamete (Sinden, 2015). This reproductive asymmetry favors an overproduction of female gametocytes to maximize fertilization success in the mosquito. However, the degree of female bias observed in peripheral blood varies and is influenced by multiple factors, including the parasite’s inbreeding (Reece et al., 2008b; Schneider et al., 2019; West et al., 2002) and the total density of circulating gametocytes (West et al., 2002).

A particularly notable result is the discrepancy between gametocyte sex ratios estimated by microscopy and those obtained with molecular markers. Molecular data consistently indicated a higher proportion of male gametocytes (average sex ratio 0.53) compared to microscopy (0.33). This may reflect a differential detectability of male and female gametocytes under the microscope, or a bias in staining or morphological identification. Alternatively, it may point to differences in the expression levels or stability of the male and female markers used. Variability in transcript abundance between male- and female-specific genes could lead to an overestimation of male gametocytes if the selected marker is more abundantly expressed or has a longer transcript half-life. Nevertheless, transcript-based methods remain a valid and powerful tool for comparing sex ratios of a given lineage between individual hosts and for monitoring changes within individuals over time.

The ability to quantify and sex *P. relictum* gametocytes with molecular markers represents a major advancement for avian malaria research. These tools will facilitate more accurate assessments of transmission potential in wild bird populations, improving our understanding of host-parasite interactions and disease dynamics. Additionally, they provide a foundation for investigating how environmental factors, such as climate change and habitat disruption, influence parasite transmission strategies.

## Acknowledgements

This work was supported by the French National Research Agency (ANR) under grant numbers ANR-23-CE02-0013 to AR, ANR-19-CE15-0007 to AT and ANR-17-CE35-0012 to SG.

## SUPPLEMENTARY TABLES

**Table S1.** Reverse transcription reagents used.

| Reagent | (μl) |
| --- | --- |
| Nuclease-free water (Ambion) | 0,29 |
| TSO 100 μM, 5AAGCAGTGGTATCAACGCAGAGTACATrGrG+G-3'(Exiqon) | 0.1 |
| 1 mM MgCl<sub>2</sub> (Ambion) | 0.06 |
| 5M Betaine solution (Affymetrix) | 2 |
| Smartscribe 5x RT Buffer (Clontech) | 2 |
| 1 mM DTT (Clontech) | 0.05 |
| 20u/μl SuperRNAsin | 0.25 |
| Smartscribe (Clontech) | 0.5 |

**Table S2.**
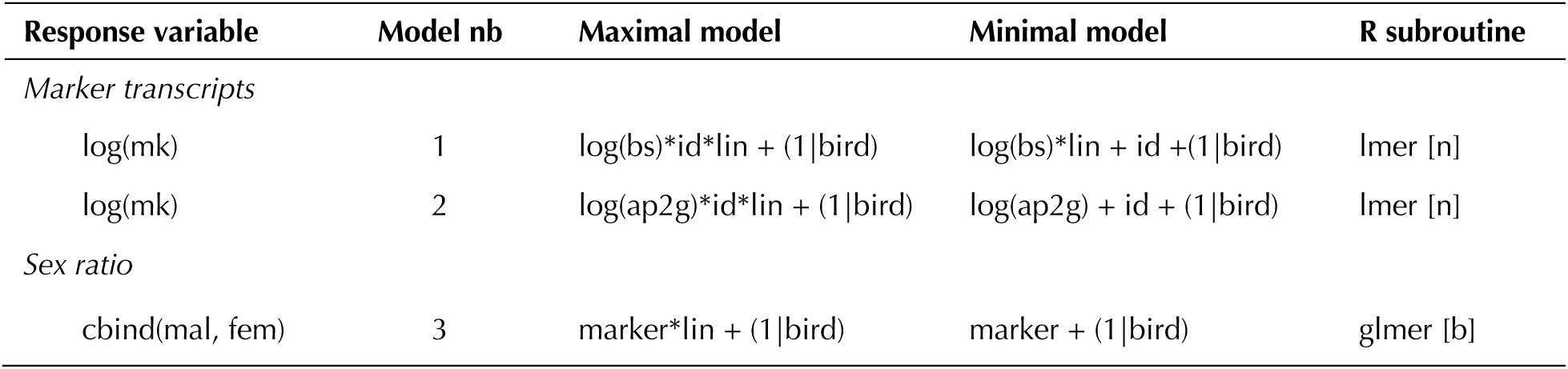
Description of the statistical models used to analyze the data. "Maximal model" represents the complete set of explanatory variables (and their interactions) included in the model. "Minimal model" represents the model containing only the significant variables and their interactions. Round brackets indicate that the variable was fitted as a random factor. Square brackets indicate the error structure used (n: normal errors, b: binomial errors). mk = number of copies of marker, bs = number of parasites in blood smear, ap2g = number of AP2-G copies, id = marker identity, lin = lineage (pSGS1, pDELURB4), bird = bird identity.

## SUPPLEMENTARY FIGURE

**Figure S1.**
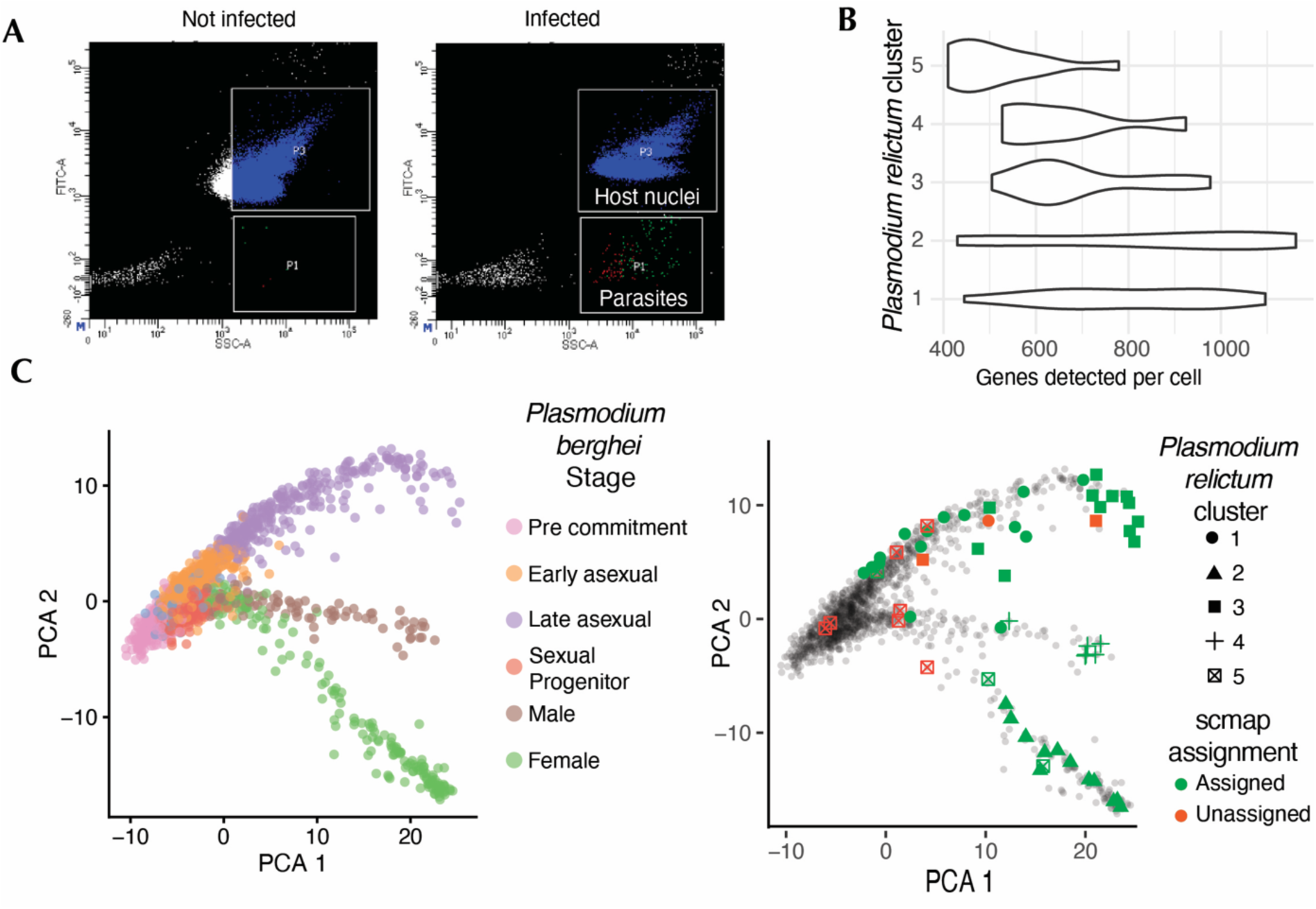
**(A)** Cytogram of non-infected and infected blood samples stained with SYBR-Green following selective lysis of host cells. Host nuclei can be seen in gate P3, while parasites are separated and sorted based on SYBR-green fluoresence and appeare in the P4 gate. **(B)** Number of genes per cell in the different clusters. **(C)** scmap analysis of the *P. relictum* transcriptomes transformed by orthology to *P. berghei* and mapped onto a reference dataset of asexual and sexual *P. berghei* stages (PMID: 36634679). Native clusters from the reference dataset are displayed on the left panel. *P. relictum* cells are shown on the right panel, centered around the top 3 reference cells to which they mapped most closely. A similarity threshold of 0.4 was applied, with cells meeting this threshold shown in green.

